# Sex-specific changes in the aphid DNA methylation landscape

**DOI:** 10.1101/286302

**Authors:** Thomas C. Mathers, Sam T. Mugford, Lawrence Percival-Alwyn, Yazhou Chen, Gemy Kaithakottil, David Swarbreck, Saskia A. Hogenhout, Cock van Oosterhout

## Abstract

Aphids present an ideal system to study epigenetics as they can produce diverse, but genetically identical, morphs in response to environmental stimuli. Here, using whole genome bisulphite sequencing and transcriptome sequencing of the green peach aphid (*Myzus persicae*), we present the first detailed analysis of cytosine methylation in an aphid and investigate differences in the methylation and transcriptional landscapes of male and asexual female morphs. We find that methylation primarily occurs in a CG dinucleotide (CpG) context and that exons are highly enriched for methylated CpGs, particularly at the 3’ end of genes. Methylation is positively associated with gene expression, and methylated genes are more stably expressed than un-methylated genes. Male and asexual female morphs have distinct methylation profiles. Strikingly, these profiles are divergent between the sex chromosome and the autosomes; autosomal genes are hypo-methylated in males compared to asexual females, whereas genes belonging to the sex chromosome, which is haploid in males, are hyper-methylated. Overall, we find correlated changes in methylation and gene expression between males and asexual females, and this correlation is particularly strong for genes located on the sex chromosome. Our results suggest that differential methylation of sex-biased genes plays a role in *M. persicae* sexual differentiation.

## Introduction

Cytosine methylation is an epigenetic mark found in many eukaryotic organisms (Bewick et al. 2016, 2017; Feng et al. 2010; Zemach and Zilberman 2010). In mammals, cytosine methylation mainly occurs in a CG dinucleotide context (CpG) (Suzuki and Bird 2008). However, in human embryonic stem cells (Guo et al. 2014), and human and mouse oocytes (Guo et al. 2014; Okae et al. 2014), cytosines are methylated in other sequence contexts (non-CpG). Plants also have high levels of non-CpG methylation that is maintained by a set of specialised CHROMOMETHYLASE enzymes not found in other eukaryotes (Bewick et al. 2017). CpG methylation is extensively detected throughout mammalian and plant genomes and often associated with suppression of gene, or transposable element, expression. In contrast, insect genomes have sparse cytosine methylation that is almost exclusively restricted to CpG sites in gene bodies (Zemach et al. 2010). Furthermore, rather than supressing gene expression, insect CpG methylation is associated with high and stable gene expression (Wang et al. 2013; Patalano et al. 2015; Xiang et al. 2010; Libbrecht et al. 2016).

Social Hymenoptera have been used as a model system to study the function of insect DNA methylation and its role caste determination (Yan et al. 2014). However, replicated experimental designs have recently shown high between-sample variation (low repeatability) and no evidence of statistically significant differences in CpG methylation between social insect castes (Libbrecht et al. 2016). Furthermore, DNA methylation has a patchy distribution across the insect phylogeny, having been lost in many species, and appears to be dispensable for the evolution of sociality (Bewick et al. 2016). Development of additional model systems is therefore desirable to gain a deeper understanding of the function of cytosine methylation in insects.

Aphids have a functional DNA methylation system (Bewick et al. 2016; Hunt et al. 2010; Walsh et al. 2010) and are an outgroup to holometabolous insects (Misof et al. 2014), which have been the main focus of research into insect DNA methylation to date. Furthermore, aphids display extraordinary phenotypic plasticity during their life cycle (Dixon 1977), in the absence of confounding genetic variation, making them ideal for studying epigenetics (Srinivasan and Brisson 2012). During the summer months, aphids are primarily found as alate, asexually reproducing, females. These asexual females are able to produce morphologically distinct morphs in response to environmental stimuli. This can include the induction of winged individuals in response to crowding (Müller et al. 2001), or the production of sexually reproducing forms in response to changes in temperature and day length (Blackman 1971b). In the case of the production of sexually reproducing individuals, sex is determined by an XO chromosomal system, where males are genetically identical to their mothers apart from the random loss of one copy of the X chromosome (Wilson et al. 1997). Differences between aphid morphs are known to be associated with large changes in gene expression (Jaquiéry et al. 2013; Purandare et al. 2014), but whether or not changes in cytosine methylation are also involved is unknown.

Here, we performed the first in-depth, genome-wide, analysis of aphid DNA methylation. We find that asexual females and males have distinct expression and methylation profiles and that changes in methylation differ between the X chromosome and autosomes. In males, the autosomes are hypo-methylated relative to asexual females whilst the X chromosome is hyper-methylated. Changes in gene expression and methylation between asexual females and males are correlated, and this correlation is strongest for X-linked genes. Taken together our findings suggest a role for DNA methylation in the regulation of aphid gene expression and the establishment of sexual dimorphism.

## Results and Discussion

### Extensive sex-biased expression between asexual females and males

To identify genes with sex-biased expression in *M. persicae* clone O, we sequenced the transcriptomes of asexual females and males (six biological replicates each) using RNA-seq (**Supplementary Table 1**). After mapping these reads to the *M. persicae* clone O genome (Mathers et al. 2017), we conducted differential expression analysis with edgeR (Robinson et al. 2009). Genes were classified based on whether their expression was significantly biased (edgeR; Benjamini-Hochberg (BH) corrected *p* < 0.05 and absolute fold change (FC) > 1.5) towards asexual females (FAB genes) or males (MB genes). We also considered the magnitude of sex bias, classifying genes as either moderately sex-biased (1.5 ≤ FC < 10, for FAB or MB) or extremely sex-biased (FC ≥ 10, for FAB+ or MB+). In total, 3,433 genes exhibited sex-biased expression (**Figure 1a**; **Supplementary Table 2**), representing 19 % of all annotated *M. persicae* genes and 33 % of all genes with detectable expression (> 2 counts-per-million in at least 3 samples, n = 10,334). MB genes outnumbered FAB genes by 18 % (1,778 vs 1,505, binomial test; p = 1.02 × 10^-6^) and only a handful of FAB+ genes (15) were observed compared to 135 MB+ genes (binomial test; *p* = 1.28 × 10^-25^; **Figure 1b**). This is consistent with patterns of sex-biased gene expression in the pea aphid (Purandare et al. 2014) and other invertebrates such as *Caenorhabditis* (Thomas et al. 2012; Albritton et al. 2014) and *Drosophila* (Zhang et al. 2007), which also show a tendency towards an excess of male-biased genes.

**Figure 1.**
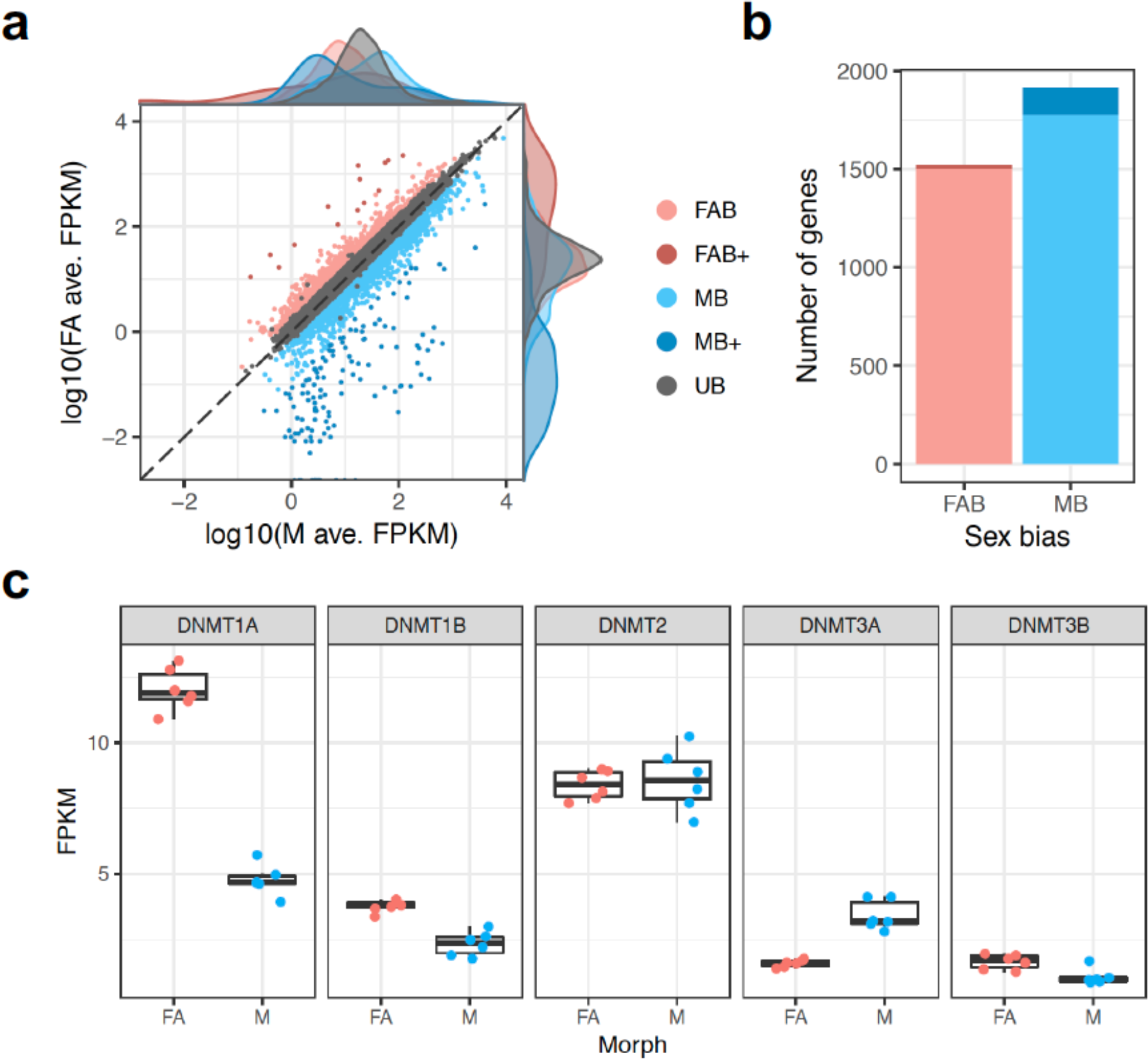
Differential gene expression between *M. persicae* asexual females and males. (**a**) Male (M; x-axis) and asexual female (FA; y-axis) gene expression expressed as log_10_ fragments per kilobase of transcript per million mapped reads (FPKM) averaged over 6 biological replicates for genes retained for differential expression (DE) analysis with edgeR (n = 10,334). DE genes are coloured according to the direction and magnitude of sex-bias (see main text). UB = unbiased expression (edgeR; Benjamini-Hochberg (BH) corrected *p* > 0.05 and absolute fold change (FC) > 1.5). (**b**) Male-biased (MB) genes significantly outnumber asexual female-biased (FAB) genes. (**c**) Asexual females and males differ significantly in expression at two out of five DNA methyltransferase genes (DNMT1a and DNMT3a; edgeR; BH corrected *p* < 0.05 and FC > 1.5). DNMT1b and DNMT3b are also significantly down-regulated in males (edgeR; BH corrected *p* = 6.35 x 10^-6^ and 0.039, respectively). However, the absolute FC of these genes falls below our cut-off of absolute FC > 1.5 (FC = 1.42 and 1.35, respectively).

### Methylation genes are differentially expressed

Next, we used our transcriptome data to investigate expression patterns of known methylation genes in *M. persicae* asexual females and males. Genome-wide patterns of DNA methylation in animals are maintained by a toolkit of DNA methyltransferase genes (Schübeler 2015). *De novo* DNA methylation is established by DNMT3 and DNA methylation patterns are maintained by DNMT1 (Law and Jacobsen 2010). An additional homolog of DNMT1 and DNMT3, DNMT2, is responsible for tRNA methylation (Goll et al. 2006) and not involved in DNA methylation. Conservation of the DNA methylation toolkit varies across insects (Bewick et al. 2016) with DNMT1 being associated with the presence of detectable levels of DNA methylation. Aphid genomes contain a full complement of DNA methylation genes with two copies of DNMT1, a single copy of DNMT2, and two copies of DNMT3 (Mathers et al. 2017; Nicholson et al. 2015; Walsh et al. 2010). We find that DNMT1a is down-regulated in males, relative to asexual females (edgeR; BH corrected *p* = 5.84 x 10^-40^, abs. FC = 2.25), and DNMT3a is up-regulated in males (edgeR; BH corrected *p* = 3.18 × 10^-14^, abs. FC = 2.44) (**Figure 1c**). DNMT1b and DNMT3b are also down-regulated in males (edgeR; BH corrected *p* = 6.35 × 10^-6^ and 0.039, respectively), however the FC of these genes falls below our 1.5-fold threshold. In contrast, the tRNA methyltransferase DNMT2 is uniformly expressed (edgeR; BH corrected *p* = 0.067). These results suggest that changes in DNA methylation may be involved in the establishment of sexual dimorphism in *M. persicae*.

### Genome-wide methylation patterns in *M. persicae*

DNA methylation has been poorly studied in insects outside of Holometabola and has only been superficially described in Hemiptera as part of a broad scale comparative analysis (Bewick et al. 2016). We therefore first sought to characterise genome-wide patterns of methylation in *M. persicae* before going on to investigate sex-specific changes in DNA methylation levels between asexual female and male morphs. To characterise genome-wide DNA methylation levels at base-level resolution, we sequenced bisulphite-treated DNA extracted from whole bodies of asexual females and males (three biological replicates each) derived from the same clonally reproducing population (clone O), and mapped these reads to the *M. persicae* clone O genome (Mathers et al. 2017) using Bismark (Krueger and Andrews 2011). After removal of ambiguously mapped reads and PCR duplicates, each replicate was sequenced to between 24x and 37x average read depth (**Supplementary Table 3**), resulting in 7,836,993 CpG sites covered by at least 5 reads in all samples.

*M. persicae* individuals harbour an obligate endosymbiont, *Buchnera aphidicola*, which lacks a functional DNA methylation system (van Ham et al. 2003). We made use of *Buchnera* derived reads in each sample to establish rates of false positive methylation calls caused by incomplete cytosine conversions by mapping each sample to the *M. persicae Buchnera* genome (Mathers et al. 2017) and quantifying methylation levels (**Supplementary Table 4**). The average methylation level in *Buchnera* for Cs in any sequence context was 0.45% ± 0.68 (mean ± SD), indicating that bisulphite treatment of the aphid DNA was 99.55% efficient and was consistent across samples. Based on this, we assessed methylation levels in *M. persicae* for C’s in a CpG, CHH and CHG context. Only Cs in a CpG context had methylation levels higher than the false positive rate in *B. aphidicola*, indicating that CpG methylation is the predominant form of DNA methylation in *M. persicae* (**Figure 2a**). Overall, global CpG methylation levels (2.93% ± 0.32% of Cs; mean ± SD) were similar to those reported in other hemipteran insects (2 – 4 %) and higher than in Hymenoptera (0.1 – 2.2 %) (Bewick et al. 2016). Exons were highly enriched for methylated CpGs relative to the rest of the genome (χ^2^ = 1.07 × 10^8^, d.f. = 1, *p* < 2.2 × 10^-16^), with only 7.7% of methylated CpGs occurring in intergenic regions (**Figure 2b and c**). Identification of significantly methylated CpG sites using a binomial test that incorporates the false positive methylation rate (derived from *Buchnera*) showed that methylation is non-randomly distributed across *M. persicae* gene bodies, with methylated CpG sites biased towards the 3’ end of genes despite the total number of CpG sites being much higher at the 5’ ends of genes (**Figure 2d**). This is likely driven by high rates of methylation in 3’ UTRs (**Figure 2c**). Interestingly, methylation bias towards the 3’ end of genes is potentially a unique feature of aphid or hemipteran CpG methylation as insects from other orders show an opposite bias, with higher methylation at the 5’ end of genes (Bonasio et al. 2012; Hunt et al. 2013; Zemach et al. 2010).

**Figure 2.**
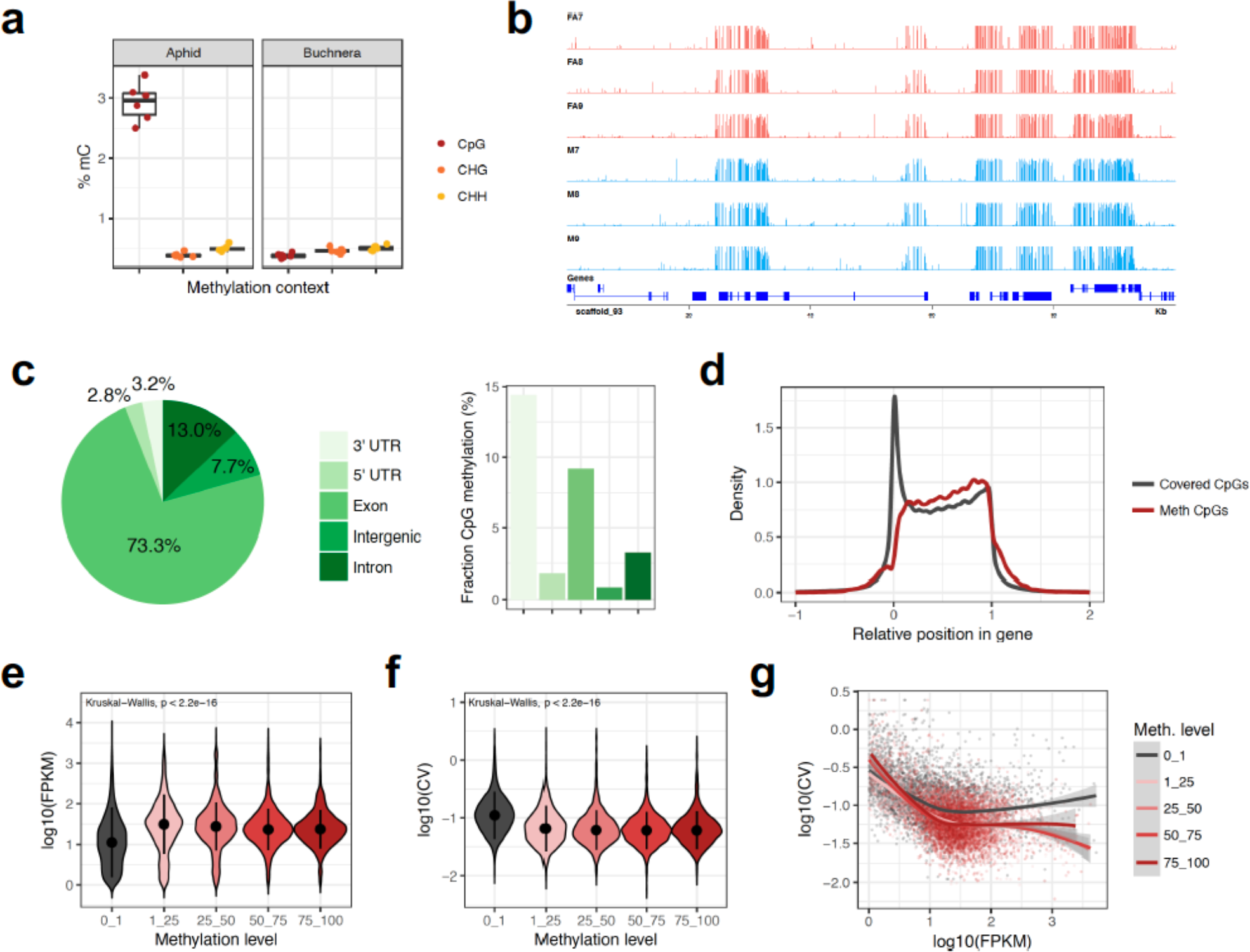
The *M. persicae* methylome. (**a**) *Boxplots* showing the proportion of methylated cytosines (mC) by sequence context (CpG, CHG and CHH) for *M. persicae* and its obligate endosymbiont *Buchnera aphidicola*, which lacks a functional methylation system. (**b**) Example genome browser view showing the distribution of CpG methylation in asexual females and males across the first 100Kb of scaffold_93. (**c**) The distribution of methylated CpGs across genomic features and the proportion of methylated CpGs in each feature. Methylated and un-methylated CpG counts were summed across all replicates. (**d**) The distribution of all covered CpG sites (min. 5 reads per sample) and significantly methylated CpG sites (*binomial test*, BH-corrected p < 0.05) across *M. persicae* gene bodies. −1 = 1Kb upstream, 0 = transcription start site, 1 = transcription stop site, 2 = 1Kb upstream. (**e**) The distribution of RNA-seq expression levels in asexual females (log_10_ FPKM) for un-methylated (0 - 1% CpG methylation) and methylated genes (FPKM = Fragments Per Kilobase of transcript per Million). Expression values were averaged across six biological replicates and methylation levels averaged across three biological replicates. Only genes with average expression levels of at least 1 FPKM in males and asexual females were included. Dots and whiskers inside the *violin plots* indicate median and interquartile range respectively. (**f**) As for (**e**) but showing the distribution of variation in expression between the six asexual female RNA-seq replicates (measured as the log_10_ transformed coefficient of variation (log_10_ CV) of FPKM) for un-methylated (0 - 1% CpG methylation) and methylated genes. (**g**) The relationship between the mean and the CV of gene expression for unmethylated and methylated genes with a trend line for each methylation level shown as a LOESS-smoothed line with shaded areas indicating the 95% CI. The difference between the grey (unmethylated; 0 - 1% CpG methylation) and pink/red lines (methylated; > 1% CpG methylation) shows that methylation reduces the between-replicate variation in gene expression, particularly in highly expressed genes. The negative correlation and downwards slope of trend lines shows that higher expressed genes are better canalized, showing less between-individual variation in gene expression.

Next, we investigated the relationship between genome-wide patterns of DNA methylation and gene expression using data for asexual females (**Supplementary Table 5**). We find that the presence DNA methylation is positively associated with gene expression, with methylated genes having significantly higher expression than un-methylated genes (**Figure 2e**). We also find that methylated genes are more stably expressed than un-methylated genes (**Figure 2f**), even after accounting for the higher expression of methylated genes (**Figure 2g**). The same patterns were also observed using male methylation and gene expression data (**Supplementary Figure 1**). Taken together, these data suggest that DNA methylation in aphids may be involved in establishing and stabilising high gene expression, as has been suggested in corals (Liew et al. 2017) and holometabolous insects (Wang et al. 2013; Patalano et al. 2015; Xiang et al. 2010; Libbrecht et al. 2016).

### Asexual females and males have distinct methylation profiles

To gain an overview of methylation differences between asexual female and male *M. persicae* morphs, we conducted principle component analysis based on methylation levels of 350,782 CpG sites significantly methylated (*binomial test*, BH-corrected *p* < 0.05) in at least one sample. Male and asexual female morphs clearly form distinct clusters, indicating reproducible differences in global CpG methylation (**Figure 3a**). To further characterise methylation differences between asexual females and males we conducted site-wise differential methylation (DM) analysis, identifying 20,964 DM CpG sites (> 15% methylation difference, BH corrected *p* < 0.05; **Supplementary Table 6**), 79% of which show a reduction in methylation (hypo-methylation) in males relative to asexual females and 21% the opposite (**Figure 3b**). This is significantly higher than expected by chance (see **Supplementary Figure 2**), and indicates that differences in methylation between asexual female and male morphs are unlikely to be due to random between-sample variation. Rather, alterations in CpG methylation appear to underpin differentiation between sexual morphs in aphids. These findings are striking given the absence of significant levels of sex-biased or caste-biased methylation in other insect systems (Libbrecht et al. 2016; Patalano et al. 2015; Herb et al. 2012; Wang et al. 2015).

**Figure 3.**
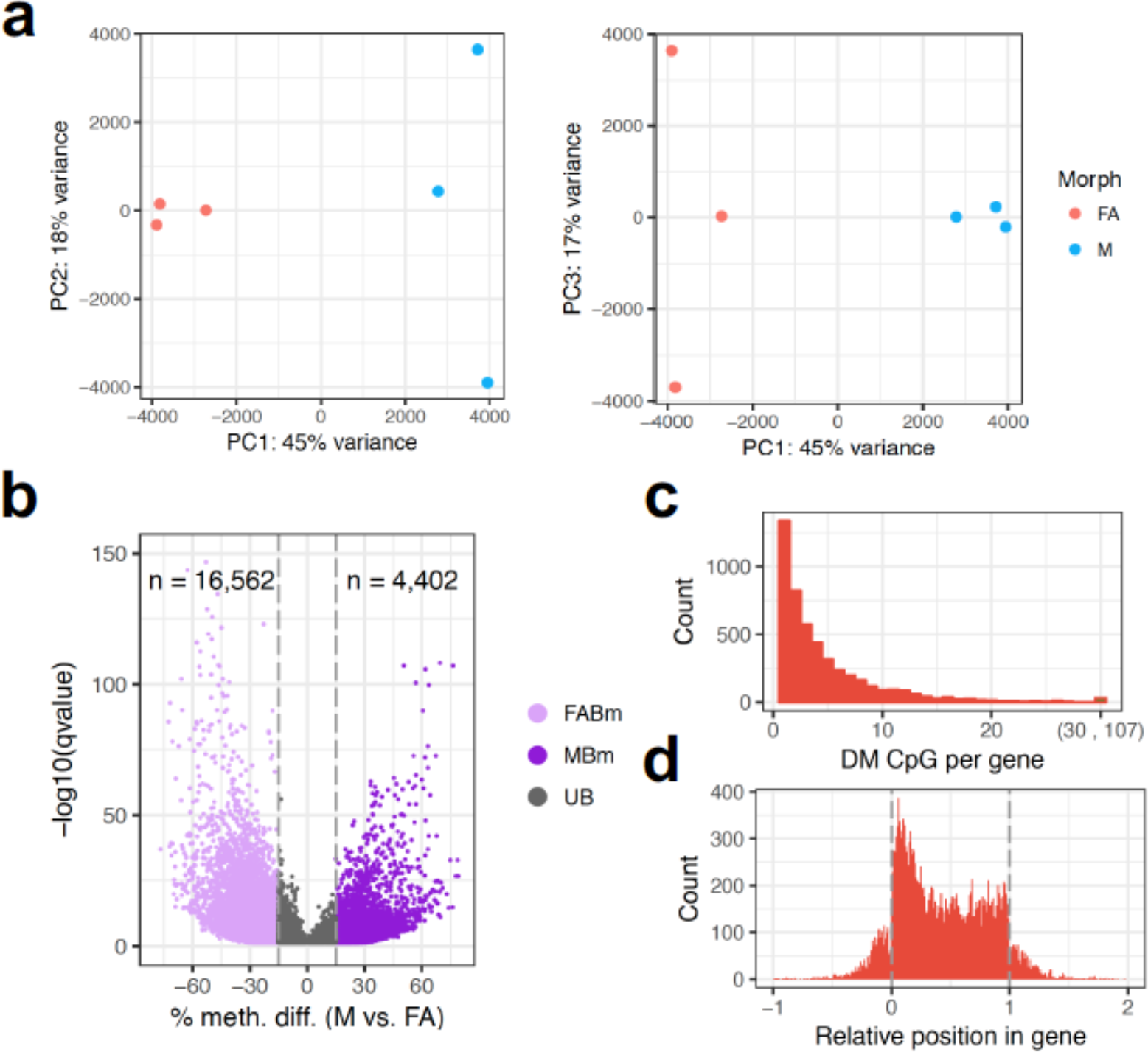
Differential methylation between *M. persicae* asexual female and male morphs. (**a**) Principle component analysis (PCA) based on methylation levels at 350,782 CpG sites significantly methylated in at least one sample. PC1 seperates the samples based on sex (45% of the variation), whilst PC2 and PC3 seperate male and asexual female replicates, respectively (explaining 18 to 17 % of the variation). (**b**) *Volcano plot* showing results of MethylKit (Akalin et al. 2012) site-wise tests of differential methylation between asexual females (FA) and males (M). Methylation differences are shown for M relative to FA. Only CpG sites showing significant differential methylation (DM) (BH corrected *p* < 0.05) are shown. A minimum methylation difference threshold of 15% per site was applied to define a site DM between FA and M. (**c**) The number of differentially methylated sites per gene (±1Kb flanking region). (**d**) The distribution of DM CpG sites along *M. persicae* gene bodies. −1 = 1Kb upstream, 0 = transcription start site, 1 = transcription stop site, 2 = 1Kb upstream.

Overlap analysis revealed that the majority (92%) of DM CpG sites between asexual females and males were located in gene bodies (± 1 Kb), with genes having between 1 and 107 DM CpG sites (**Figure 3c**). These DM CpG sites were non-randomly distributed along gene bodies, being biased towards the 5’ end of genes (**Figure 3d**). As such, whilst overall methylation levels are biased towards the 3’ end of genes, sites with variable methylation are more likely to be at the 5’ end. To directly correlate gene body methylation levels with gene expression, we also performed DM analysis at the gene level (**Supplementary Table 7**). This identified 1,344 DM genes with > 10% methylation difference (BH corrected *p* < 0.05), of which 205 showed significant male-biased methylation and 1,129 asexual female-biased methylation (**Figure 4a and b**). Considering genes with variable methylation, males have undergone a global loss of gene body methylation relative to asexual females (Wilcoxon signed-rank test, *p* < 2.2 × 10^-22^; **Figure 4c**).

**Figure 4.**
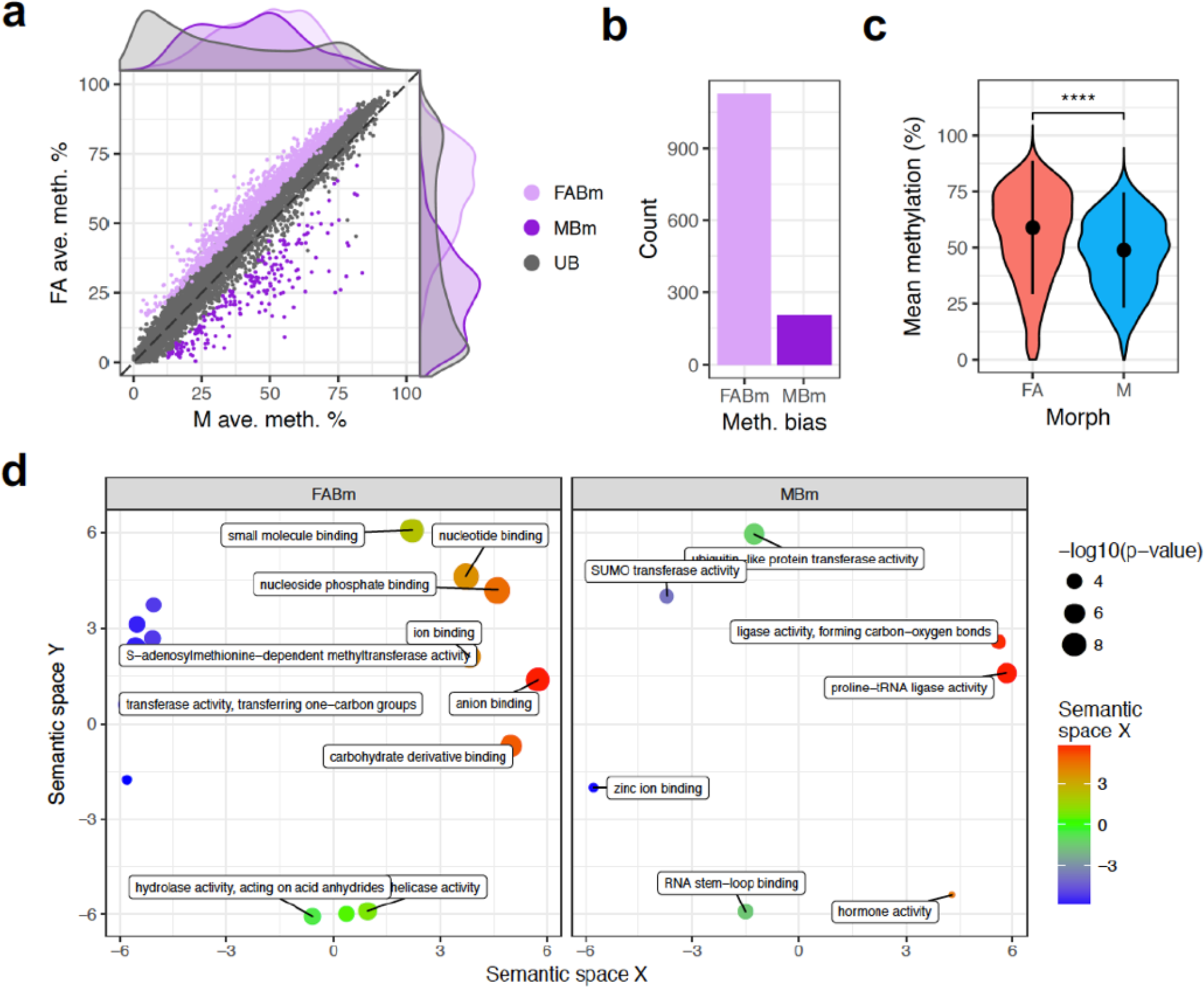
Genome-wide changes in gene body methylation between asexual female and male morphs. (**a**) Male (M; x-axis) and asexual female (FA; y-axis) gene-wise methylation levels averaged over 3 biological replicates for genes methylated > 1% in at least one of the two morphs (n = 6,699). Differentially methylated (DM) genes (MethylKit; > 10% methylation difference, BH corrected *p* < 0.05) are coloured according to the direction of sex-bias: MBm = male biased methylation, FABm = female-biased methylation, UB = unbiased methylation. (**b**) FABm genes outnumber MBm genes. (**c**) *Violin plot* showing the distribution of mean methylation level in FA and M for DM genes. Dots and whiskers indicate median and interquartile range respectively; **** = Wilcoxon signed-rank test *p* < 0.0001. (**d**) Enriched GO terms relating to molecular function plotted in semantic space for FABm genes and MBm genes (for terms relating to biological process see **Supplementary Figure 3**). A full list of enriched GO terms for each DM class and functional category is given in **Supplementary Table 8**).

Gene ontology (GO) term enrichment analysis showed that asexual female-biased methylation and male-biased methylation genes were both enriched for GO terms relating to core biological processes, including metabolism and regulation of gene expression (**Figure 4d**; **Supplementary Figure 3**; **Supplementary Table 8**). Protein SUMOylation is enriched among genes with male-biased methylation. This is interesting because protein SUMOylation is essential for dosage compensation of the *C. elegans* sex chromosome (Pferdehirt and Meyer 2013) and plays a key role in insect development and metamorphosis (Ureña et al. 2015). Changes in methylation may therefore underpin core processes involved in aphid polyphenism and sex determination. Consistent with this, we also find enrichment of hormone signalling amongst genes with male-biased methylation, with 3 insulin genes hyper-methylated in males (2 not expressed, 1 has male-specific expression). Insulin receptors determine alternative wing morphs in planthoppers (Xu et al. 2015) and have been shown to interact with the core sex determination gene TRANSFORMER-*2* (Zhuo et al. 2017).

### The X chromosome has distinct patterns of expression and methylation

We identified X-linked scaffolds in the *M. persicae* genome assembly based on the ratio of male to asexual female bisulphite sequencing coverage. This approach takes advantage of the hemizygous condition of the X chromosome in males, which should result in X-linked scaffolds having half the read depth of autosomal scaffolds (Jaquiéry et al. 2017). As expected, we observe a bimodal distribution in the ratio of male to asexual female scaffold coverage, with the lower coverage peak falling at approximately half the relative coverage of the higher coverage peak (**Figure 5a**; **Supplementary Table 9**). Scaffolds in this lower coverage peak are putatively derived from the X chromosome. To validate the coverage results, we mapped known X-linked (n=4) and autosomal (n=8) microsatellite loci to the clone O genome and retrieved the male to asexual female coverage ratios of their corresponding scaffolds. The coverage of these known sex-linked scaffolds also exactly matches expectations (**Figure 5a**; **Supplementary Table 10**). Using a cut-off of adjusted coverage, we identified 748 X-linked scaffolds and 1,852 autosomal scaffolds, totalling 68.7 and 239.7 Mb of sequence respectively (**Supplementary Figure 4**). Scaffolds assigned to the X chromosome therefore account for 22.3% of the assembled (scaffolds ≥ 20Kb) *M. persicae* clone O genome. This is in line with expectations given the most common *M. persicae* karyotype of 2n = 12 and that the X chromosome is larger than the autosomes (Blackman 1971a). Using the chromosomal assignment of scaffolds, we were able to assign 3,110 gene models to the X chromosome and 10,768 to autosomes, leaving 4,555 (24.7%) gene models on unassigned scaffolds shorter than 20 Kb. The number of identified X-linked genes is not different to expectations based on the assembled size of the respective chromosomal regions (binomial test, *p* = 0.65). However, we find that the X chromosome is depleted in coding sequence (CDS) compared to the autosomes (6.3% vs 6.5%; χ^2^ = 5,821.5, d.f. = 1, *p* < 2.2 × 10^-16^). This is due to the reduced CDS length of X-linked genes (Wilcoxon signed-rank test, *p* = 4.2 × 10^-4^; **Supplementary Figure 5**).

**Figure 5.**
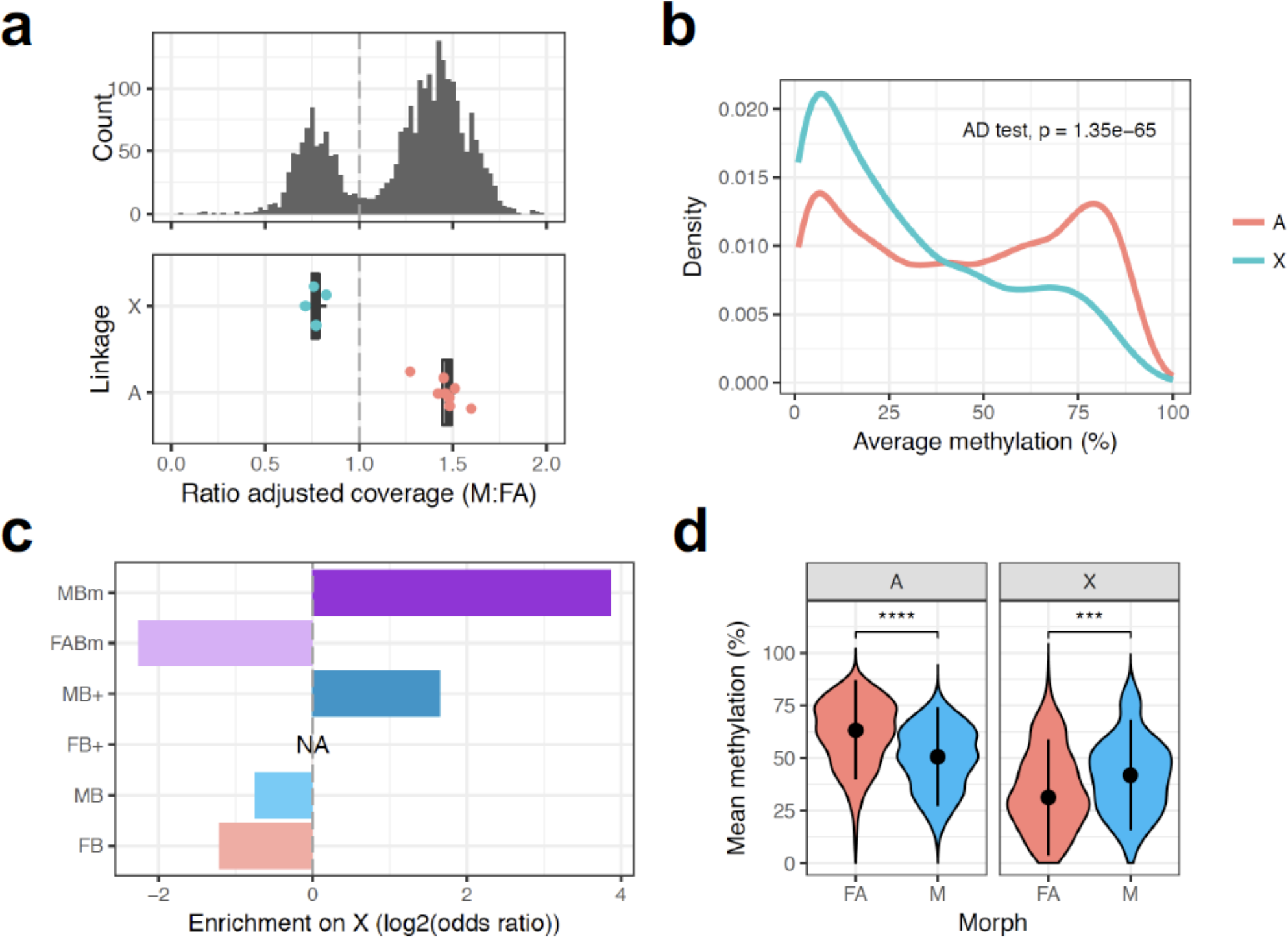
Distinct patterns of methylation and expression between the *M. persicae* X chromosome and autosomes. (**a**) X-linked and autosomal scaffolds (≥ 20Kb) in the *M. persicae* genome were identified based on the relative coverage of BS-seq reads in males (M) compared to asexual females (FA). Given the XO sex determination system of aphids, X-linked scaffolds are predicted to have half autosomal coverage in males. A bimodal distribution in the ratio of M to FA coverage is clearly visible (upper panel). We considered scaffolds falling in the lower coverage peak (ratio of adjusted coverage < 1) as X-linked and scaffolds in the second, higher coverage peak (ratio of adjusted coverage > 1), as autosomal. The assignment of scaffolds to the X chromosome or autosomes was validated by comparing the M : FA ratio of coverage for scaffolds containing microsatellite markers on the X-chromosome (blue dots) and autosome (red dots) (lower panel). (**b**) The distribution of gene body methylation levels for X-linked and autosomal genes. (**c**) Observed / expected (odds ratio) counts of DM and DE genes on the X chromosome by expression or methylation bias category. The X chromosome is significantly enriched for MB+ genes (≥ 10-fold upregulation in M) and genes with male-biased methylation (MBm). (**d**) The distribution of mean methylation levels in asexual females (FA) and males (M) for X-linked and autosomal DM genes (MethylKit; > 10% methylation difference, BH corrected *p* < 0.05). Methylation levels are significantly higher in FA than M for autosomal genes, whereas M have a higher methylation than FA in X-linked genes (**d**) dots and whiskers inside the *violin plots* indicate median and interquartile range respectively; *** = Wilcoxon signed-rank test *p* < 0.001, **** = *p* < 0.0001.

Strikingly, the X chromosome has a distinct methylation landscape compared to the autosomes (Anderson-Darling k-sample test, *p* = 1.35 × 10^-65^; **Figure 5b**), with fewer highly methylated genes. We also find opposing dynamics of sex-biased methylation between the X chromosome and the autosomes. The X chromosome is significantly enriched for genes with male-biased methylation and depleted for genes with female-biased methylation (χ^2^ = 176.65, d.f. = 2, *p* < 2.2 × 10^-16^; **Figure 5c**). Overall, X chromosome genes are hyper-methylated in males (Wilcoxon signed-rank test, *p* = 8.6 × 10^-4^; **Figure 5d**) compared to the genome-wide pattern of hypo-methylation (Wilcoxon signed-rank test, *p* < 2.2 × 10^-16^). Mirroring differences in methylation between the X chromosome and the autosomes, we also find that the X chromosome is enriched for genes with extreme male-biased expression (χ^2^ = 42.38, d.f. = 1, *p* = 7.5 × 10^-11^; **Figure 5c**), a phenomenon also observed in the pea aphid (Jaquiéry et al. 2013). Male-biased expression of X-linked genes is therefore conserved across two distantly related aphid species, and, at least in the case of *M. persicae*, this also extends to patterns of DNA methylation.

Finally, we investigated whether changes in methylation between *M. persicae* asexual females and males are associated with changes in gene expression. The relationship between gene expression and gene body methylation is an open question in invertebrates and few studies have directly tested for changes in expression and methylation. We find that DM genes are significantly enriched for DE (χ^2^ = 7.84, d.f.= 1, *p* = 0.005), suggesting that methylation changes may be involved in the regulation of at least a subset of sex-biased genes. In support of this, we find a weak but significant positive correlation between changes in gene expression and methylation between asexual females and males when considering genes methylated (> 1%) and expressed (> 1 FPKM) in at least one of the sexes (n = 6,699; Spearman’s ρ = 0.089, *p* = 2.7 × 10^-13^; **Figure 6a**). Interestingly, this correlation is driven by X-linked genes which show a significantly stronger correlation between changes in expression and methylation than autosomal genes (GLM: F_1,6185_ = 93.07, *p* < 0.0001; **Figure 6b**). Combined with recent results demonstrating a role for chromatin accessibility in the sex-specific regulation of genes on the X chromosome and dosage compensation in the pea aphid (Richard et al. 2017), our findings suggest a key role for epigenetics in establishing patterns of X-linked gene expression in aphids.

**Figure 6.**
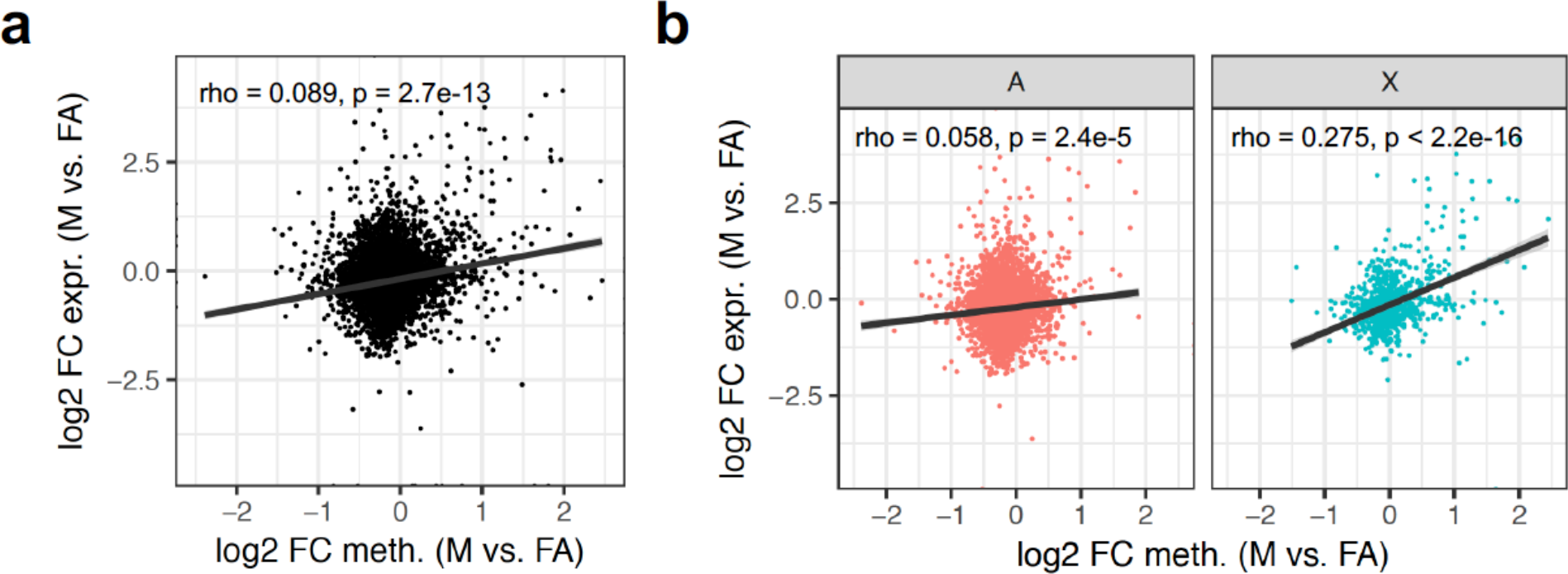
Correlated changes in expression and methylation between asexual females and males. (**a**) *Scatter plot* showing the relationship between fold-change (FC) in gene expression and methylation between asexual females (FA) and males (M) for genes expressed (> 1 FPKM) and methylated (> 1%) in at least one of the sexes (n = 6,699). Methylation levels of genes were estimated across the whole gene body and averaged across replicates. Positive values indicate increased expression or methylation in males, relative to asexual females; negative values indicate increased expression or methylation in asexual females, relative to males. (**b**) The correlation between gene expression changes and methylation changes between FA and M is significantly stronger for X-linked genes (X; n = 925) than autosomal genes (A; n = 5,272). Spearman’s ρ was used to assess significance and strength of the relationship between change in expression and methylation for each set of genes. The trend lines indicate linear fit with shaded areas indicating 95% confidence intervals.

## Conclusions

We presented the first detailed analysis of genome-wide methylation patterns in an aphid, evaluating its importance for gene expression and sexual differentiation. We found that 3,433 genes (19 % of the annotated genome) were differentially expressed between the males and asexual females, and that there was a significant excess of male-biased genes. We also found evidence suggesting that methylation plays an important role in sexual differentiation of aphids, showing that DNMT1a and b are significantly downregulated in males, whereas DNMT3a is upregulated in males. CpG methylation is the predominant form of DNA methylation in *M. persicae* and, in contrast to other insects, exons were highly enriched for methylated CpGs at the 3’ end rather than the 5’ end of genes. Methylation is positively associated with gene expression, and in addition, methylated genes are more stably expressed than un-methylated genes. Methylation was significantly reduced in males compared to asexual females, yet remarkably, the X chromosome genes of males were hyper-methylated. Given that differentially methylated genes were also significantly differentially expressed between the sexes, we propose that changes in DNA methylation play a role in *M. persicae* sexual differentiation. Our findings pave the way for future functional studies of DNA methylation in aphids, and its potential role in the remarkable evolutionary potential of these insects, and their extraordinary phenotypic plasticity.

## Methods

### Aphid rearing and sample preparation

An asexual colony of *M. persicae* clone O derived from a single apterous asexual female (Mathers et al. 2017) was maintained on *Brassica rapa* plants in long-day conditions (14h light, 22° C day time, and 20° C night time, 48% relative humidity). Male morphs were induced by transferring the colony to short-day conditions (8h light, 18° C day time, and 16° C night time, 48% relative humidity) and samples were collected 2 months after transfer. Replicate samples were harvested from the same populations, with each replicate consisting of 20 adults, with apterous asexual females collected from the long-day population, and males from the short-day population. Samples were immediately frozen in liquid nitrogen prior to RNA or DNA extraction. DNA (three biological replicates per morph) was extracted using the CTAB protocol (Marzachi et al. 1998), with the addition of a proteinase K digestion step during the initial extraction. RNA (six biological replicates per morph) was extracted using the Trizol reagent according to the manufacturers’ protocol (Sigma), and further purified using the RNeasy kit with on-column DNAse treatment (Qiagen).

### Transcriptome sequencing

RNA samples were sent for sequencing at the Earlham Institute (Norwich, UK) where twelve non-orientated libraries were constructed using the TruSeq RNA protocol v2 (Illumina #15026495 Rev.F). 1ug of total RNA was enriched for mRNA using oligo(dT) beads. The RNA was then fragmented and first strand cDNA synthesised. Following end repair and adapter ligation, each library was subjected to a bead based size selection using Beckman Coulter XP beads (Beckman Coulter Inc., Brea, CA, USA) before performing PCR to enrich for fragments containing TruSeq adapter sequences. Libraries were then pooled and sequenced on the Illumina HiSeq 2000 platform (Illumina Inc., San Diego, CA, USA) (v3 chemistry; 2 x 100 bp), generating between 15 and 57 million paired-end reads per sample. RNA-seq reads have been deposited in the NCBI short read archive (SRA) under accession number PRJNA437622.

### Gene expression analysis

Raw RNA-seq reads for each sample were trimmed for low quality bases and adapter contamination with Trim Galore! v0.4.0 using default settings for paired end reads (www.bioinformatics.babraham.ac.uk/projects/trim_galore/). Gene-level expression quantification was then performed for each sample based on the *M. persicae* clone O reference genome and gene annotation (Mathers et al. 2017), using RSEM v1.2.31 (Li and Dewey 2011) with STAR v2.5.2a (Dobin et al. 2013). Average expression and the coefficient of variation was calculated per gene for asexual females and males separately using FPKM (fragments per kilobase of transcript per million) values estimated by RSEM. We also identified differentially expressed (DE) genes between asexual females and males using edgeR (Robinson et al. 2009) based on gene-level expected counts estimated by RSEM. Only genes with greater than 2 counts-per-million in at least three samples were retained for DE analysis and we considered genes DE if they had a fold-change (FC) ≥ 1.5 and *p* < 0.05 after adjusting for multiple testing using the Benjamini-Hochberg (BH) procedure (Benjamini and Hochberg 1995).

### Bisulphite sequencing

Bisulphite sequencing library construction was performed using 500 ng genomic DNA per sample with a BIOO Scientific NEXTflex^TM^ Bisulfite-Seq Kit (Bioo Scientific Corporation, Austin, TX, USA) according to the manufacturer’s instructions with the following modifications: genomic DNA was sheared to 200-400 bp with a Covaris S2 sonicator (Covaris Inc., Woburn, MA) using the following settings: duty cycle 10%, intensity 5, 200 cycles per burst for 120 seconds. The power mode was frequency sweeping, temperature 5-6°C and water level 12. Libraries either received NEXTflex^TM^ barcode #24 (GGTAGC) or #31 (CACGAT). All purified libraries were QC checked with the Bioanalyzer DNA HS assay and further quantified by Qubit dsDNA HS Assay Kit (Life Technologies, Carlsbad, CA, USA) before pooling as pairs. Pooled libraries were further quantified by qPCR using a KAPA Library Quantification Kit - Illumina/ABI Prism (Kapa Biosystems Inc., Wilmington, MA, USA) on a StepOnePlus™ Real-Time PCR System (Life Technologies, Carlsbad, CA, USA). Sequencing was performed at the Earlham Institute (Norwich, UK) on an Illumina HiSeq 2500 (Illumina Inc., San Diego, CA, USA) using paired-end sequencing (v4 chemistry; 2 × 126 bp) with a 15% PhiX spike in, clustering to 650 K/mm^2^. In total, we generated between 70 and 127 million paired-end reads per sample.

### DNA methylation analysis

Bisulphite treated reads for each sample were trimmed for low quality bases and adapter contamination using Trim Galore! v0.4.0 with default settings (www.bioinformatics.babraham.ac.uk/projects/trim_galore/). Read pairs where one or both reads were shorter than 75bp after trimming were discarded. We then mapped the trimmed reads to the *M. persicae* clone O reference genome (Mathers et al. 2017) using Bismark v0.16.1 (Krueger and Andrews 2011). Trimmed reads were also mapped to the genome of the *M. persicae* strain of the obligate aphid endosymbiont *Buchnera aphidicola* (Mathers et al. 2017) to estimate the error rate of the C to T conversion. Reads derived from PCR duplicates and that mapped to multiple locations in the genome were removed from downstream analysis. The distribution of methylation across selected scaffolds was visualised using Sushi (Phanstiel et al. 2014).

Overall levels of methylation in a CpG, CHG and CHH sequence context were estimated directly from mapped reads with Bismark (Krueger and Andrews 2011). We also characterised CpG methylation levels of features in the *M. persicae* clone O genome based on the reference annotation (Mathers et al. 2017). Average CpG methylation levels of introns, exons, 5’ UTRs, 3’ UTRs and intergenic regions were calculated with bedtools v2.25.0 (Quinlan and Hall 2010), pooling data from all replicates and counting overlapping methylated and unmethylated CpGs. We also calculated per-gene methylation levels for asexual females and males independently in the same way. To assess the genome-wide distribution of methylated CpGs, we filtered CpG sites to those covered by at least five reads in all samples and used a binomial test to identify significantly methylated sites in each sample using the C to T conversion error rate (derived from mapping to *Buchnera*) as the probability of success and corrected for multiple testing using the BH procedure (Benjamini and Hochberg 1995), setting the FDR at 5% (BH adjusted *p* < 0.05).

Methylation differences between asexual females and males were assessed using a principle component analysis (PCA) and by identifying differentially methylated (DM) sites and genes. PCA was carried out with prcomp, implemented in R v3.2.2 (R Core Team 2017), using the methylation levels of CpG sites significantly methylated in a least one sample (binomial test, BH adjusted *p* < 0.05). We identified DM sites and genes using logistic regression implemented in MethylKit (Akalin et al. 2012) which accepts input directly from Bismark. *p* values were adjusted to Q-values using the SLIM method (Wang et al. 2011) to account for multiple testing. For the site-level analysis, we discarded CpG sites covered by less than 5 reads and those that fell into the top 0.1% of coverage. We considered sites significantly DM if they had at least a 15% methylation difference at a 5% FDR (Q < 0.05). At the gene level, we discarded genes covered by less than 20 reads which fell into the top 1% of coverage, and called genes as DM if they had at least 10% methylation difference and at a 5% FDR (Q < 0.05). A less stringent percent methylation difference was used at the gene-level as the signal of DM may be diluted over the length of the gene body. To assess the rate of false positive methylation calls caused by random variation between samples we generated a null distribution of DM calls at Q < 0.05 for a range of percentage methylation difference cut-offs using all possible, non-redundant, pairs of samples where an asexual female sample is grouped with a male sample (n = 19). Overall, these random pairings resulted in a low number of DM calls, indicating our results are robust (**Supplementary Figure 2a and b**).

### X chromosome identification

We used our whole-genome bisulphite sequencing data for males and asexual females to identify X–linked scaffolds in the *M. persicae* clone O genome assembly based on the ratio of male to asexual female coverage using a procedure similar to Jaquiéry et. al (2017). BAM files generated by MethylKit were merged for each morph using Picard v2.1.1 (http://broadinstitute.github.io/picard/) to maximise the depth of coverage. We then calculated per site sequence depth with SAMtools v1.3 (Li et al. 2009). The average depth of the pooled asexual female and male samples was 79x and 90x, respectively. We then calculated the ratio of male median depth of coverage to asexual female median depth of coverage for all scaffolds longer than 20 Kb, normalising male coverage to that of asexual female coverage (multiplying male median coverage by 79 / 90). This resulted in a clear bimodal distribution with modes at ~0.75 and ~1.5 (**Figure 5a**). We applied a cut-off of male to asexual female normalised median coverage ratio < 1 to assign scaffolds to the X chromosome and > 1 to assign scaffolds to the autosomes. To validate the coverage results, we mapped known X-linked (n=4) and autosomal (n=8) microsatellite loci from Sloane et. al. (2001) and Wilson et. al. (2004) to the clone O genome with blastn and retrieved coverage ratios for their respective scaffolds.

### Testing for correlation between changes in methylation and expression

To investigate the relationship between changes in gene expression and methylation we compared expression and methylation levels of genes in males and asexual females. Using average expression (FPKM) and methylation levels, we calculated the log_2_ FC in expression (FC_Expr_) and methylation (FC_Meth_), and tested for correlation using Spearman’s ρ (rho). We also investigated the effect of chromosomal location (X chromosome vs. autosomes) on the relationship between gene expression and methylation using a general linear model (GLM). The GLM was formulated with FC_Expr_ as the response variable, and FC_Meth_ as a covariate, crossed with chromosome (as fixed factor). This interaction term tests whether the slopes of the regression lines of the X chromosome and autosomes run parallel.

### Annotation of methyltransferase genes

Amino acid sequences of human DNA methyltransferase genes were blasted against annotated protein sequences of *Myzus persicae* Clone O (Mathers et al., 2017). The top *M. persicae* clone O hit for each gene was then used to blast against the *M. persicae* protein set in an iterative fashion until no additional genes were identified. The E value were set as 1E-10.

### GO term enrichment analysis

Go term enrichment analysis of specific gene sets was performed with BINGO (Maere et al. 2005) using the complete *M. persicae* clone O proteome as the reference set. Redundant terms were then removed with REVIGO (Supek et al. 2011).

## Data availability

Raw RNA-seq and BS-seq data generated for this study have been deposited in the NCBI short read archive under accession number PRJNA437622.

## Acknowledgements

This work was supported by a BBSRC Industrial Partnership Award (BB/L002108/1) to S.H., D.S. and C.v.O. and co-funded by Syngenta, the Plant Health Institute Strategy Programme (BB/P012574/1) awarded to the John Innes Centre, Gatsby Charitable Foundation funding to S.H., a BBSRC award (BB/N02317X/1) to C.v.O., as well as support by the Earth & Life Systems Alliance (ELSA). Next-generation sequencing and library construction was delivered via the Biotechnology and Biological Sciences Research Council (BBSRC) National Capability in Genomics and Single Cell (BB/CCG1720/1) at Earlham Institute by members of the Genomics Pipelines Group. We thank Anna Jordan at the JIC insectary for assistance with aphid rearing and identification of aphid morphs.

## Author contributions

T.C.M., S.T.M., Y.C., D.S., S.H. and C.v.O. conceived the study. T.C.M. performed bioinformatics analysis with additional analysis performed by C.v.O, G.K. and Y.C.. S.T.M. performed aphid morph phenotyping and extracted DNA and RNA. L.P.A. constructed bisulphite sequencing libraries. T.C.M, C.v.O and S.H. wrote the manuscript. All authors read, edited and approved the final manuscript.

